# Quantifying chemodiversity considering biochemical and structural properties of compounds with the R package *chemodiv*

**DOI:** 10.1101/2022.06.08.495236

**Authors:** Hampus Petrén, Tobias G. Köllner, Robert R. Junker

## Abstract

- Plants produce large numbers of phytochemical compounds affecting plant physiology and interactions with their biotic and abiotic environment. Recently, chemodiversity has attracted considerable attention as an ecologically and evolutionary meaningful way to characterize the phenotype of a mixture of phytochemical compounds.
- Currently used measures of phytochemical diversity, and related measures of phytochemical dissimilarity, generally do not take structural or biosynthetic properties of compounds into account. Such properties can be indicative of the compounds’ function and inform about their biosynthetic (in)dependence, and should therefore be included in calculations of these measures.
- We introduce the R package *chemodiv*, which retrieves biochemical and structural properties of compounds from databases and provides functions for calculating and visualizing chemical diversity and dissimilarity for phytochemicals and other types of compounds. Our package enables calculations of diversity that takes the richness, relative abundance and – most importantly – structural and/or biosynthetic dissimilarity of compounds into account. We illustrate the use of the package with examples on simulated and real datasets.
- By providing the R package *chemodiv* for quantifying multiple aspects of chemodiversity, we hope to facilitate investigations of how chemodiversity varies across levels of biological organization, and its importance for the ecology and evolution of plants and other organisms.

## Introduction

Plants produce an astonishing diversity of phytochemical compounds (Kessler & Kalske, 2018; Wang *et al*., 2019). With functions such as chemical defence, attractant or repellent signalling and protection against abiotic stressors, phytochemicals (also referred to as secondary metabolites) are crucial for mediating mutualistic and antagonistic interactions between plants and other organisms and the abiotic environment (Hartmann, 2007; Junker & Tholl, 2013; Kessler & Kalske, 2018; Whitehead *et al*., 2021b). Understanding the evolutionary processes generating this phytochemical diversity, and the ecological functions of it are central goals in the field of chemical ecology (Fraenkel, 1959; Ehrlich & Raven, 1964; Hartmann, 2007; Raguso *et al*., 2015).

Traditionally, research has mostly focused on understanding the function (e.g. herbivore protection or pollinator attraction) of individual phytochemical compounds (Richards *et al*., 2016; Dyer *et al*., 2018). However, phytochemicals occur in multicompound mixtures, the composition of which represents a complex phenotype that may vary along multiple dimensions (Marion *et al*., 2015). Recently the concept of chemodiversity has received increased attention as a way to quantify this phenotype (Junker *et al*., 2018; Wetzel & Whitehead, 2020; Müller *et al*., 2020). Chemodiversity can be quantified using diversity indices (Doyle, 2009; Hilker, 2014; Marion *et al*., 2015; Kessler & Kalske, 2018; Wetzel & Whitehead, 2020), and multiple studies have found that function may be dependent on a diverse mixture of compounds (e.g. Iason *et al*., 2005; Bruce *et al*., 2005; Richards *et al*., 2015; Junker *et al*., 2018; Tewes *et al*., 2018; Whitehead *et al*., 2021a; Cosmo *et al*., 2021). Less appreciated is the fact that phytochemical compounds are produced by a limited number of biosynthetic pathways and are characterized by different chemical structures (Wink, 2010; Wang *et al*., 2019). Considering such properties of compounds as a part of the phytochemical phenotype can be important to account for interdependences due to shared biosynthetic pathways (Junker, 2018; Junker *et al*., 2018), and crucially, a factor contributing to explaining the function of phytochemicals (Wetzel & Whitehead, 2020; Cosmo *et al*., 2021).

Chemodiversity is often measured using indices such as Shannon’s diversity index. While originally used to quantify species diversity, numerous studies have used diversity indices to quantify phytochemical diversity at different levels of biological organization, and explored its effects on ecological interactions and evolutionary processes. This includes examples where phytochemical diversity influences insect performance (Tewes *et al*., 2018; Glassmire *et al*., 2020), shapes patterns of herbivory and insect diversity across plant communities (Richards *et al*., 2015; Salazar *et al*., 2016), and where it changes over evolutionary time in different plant genera (Becerra *et al*., 2009; Cacho *et al*., 2015). Mechanistically, a high diversity of compounds might be selected for and enhance function in a number of different ways (Wetzel & Whitehead, 2020). Synergistic effects may cause the effect of a mixture of compounds to be larger than the sum of the effects of individual compounds (Richards *et al*., 2016). Alternatively, a diverse set of phytochemicals may result from the multitude of interactions plants experience, each imposing selection on different compounds with different functions (Berenbaum & Zangerl, 1996; Iason *et al*., 2011; Junker, 2016). Regardless of the exact mechanism, under each scenario an increased diversity of phytochemical compounds within a plant may increase its fitness.

Using indices such as Shannon’s diversity, most studies on phytochemical diversity consider compound richness and evenness, but ignore disparity, the third component of diversity (Daly *et al*., 2018). Analogous to measures of functional diversity, where species’ traits are included in calculations of indices such as Rao’s quadratic entropy index (Petchey & Gaston, 2006), the biosynthetic and/or structural disparity of phytochemicals (hereafter referred to as compound dissimilarity) can and should be included in calculations of phytochemical diversity. All else equal, a phytochemical mixture of structurally dissimilar compounds produced by different biosynthetic pathways is arguably more diverse than a mixture of less dissimilar compounds from a single biosynthetic pathway. How dissimilar the compounds in a phytochemical mixture are, is thus a crucial component of the mixture’s overall diversity. A higher structural diversity among compounds might mediate interactions with or increase effects against a broader set of interacting organisms or influence synergies between compounds (Becerra *et al*., 2009; Whitehead *et al*., 2021a; Cosmo *et al*., 2021; Philbin *et al*., 2022), thereby affecting function. A few methods to measure such structural variation exist. Quantifications of compound dissimilarity based on tandem (MS/MS) mass spectra (Wang *et al*., 2016), have been used in metabolomics (Tripathi *et al*., 2021) and ecology (Sedio *et al*., 2017) to calculate sample dissimilarities and construct molecular networks. Additionally, Richards *et al*. (2015) pioneered quantifying phytochemical diversity using ^1^H-NMR spectra with a measure reflective of both inter-molecular and intra-molecular diversity. Such measures have been shown to influence plant-insect interactions (Richards *et al*., 2015; Sedio, 2017; Glassmire *et al*., 2019; Sedio *et al*., 2020; Cosmo *et al*., 2021; Philbin *et al*., 2022), indicating that structural diversity may be an important factor shaping ecological interactions. Phytochemical data, however, is often analysed using standard GC-MS or LC-MS methods, where individual compounds are identified and quantified. We propose using similar methods to quantify compound dissimilarity for such datasets. By quantifying compound dissimilarities for datasets with identified compounds (Box 2), and calculating phytochemical diversity and dissimilarity of samples using measures of functional Hill diversity and Generalized UniFrac dissimilarities (Chen *et al*., 2012; Chao *et al*., 2014) (Box 1), we aim to enable chemical ecologists to quantify all components, including the richness, evenness and disparity, of phytochemical diversity.

We introduce *chemodiv*, a package for analyses of chemodiversity in the statistical software R (R Core Team, 2022). The package allows users, with data on relative abundances of identified phytochemical compounds in different samples, or any other type of chemical composition data, to quantify different types of chemical diversity and dissimilarity. These calculations include all components of diversity, where the richness, evenness and, importantly, the biosynthetic and/or structural properties of the compounds are considered. By using such comprehensive measures, we hope that researchers will be able to efficiently test what dimensions of phytochemical diversity are most important in shaping interactions between plants and their biotic and abiotic environment.

### Box 1. Measures of diversity and dissimilarity

Diversity can be divided into components of richness, evenness and disparity (Daly *et al*., 2018). The most simple diversity measure is simply the richness, in this case the number of phytochemical compounds detected in a sample. Studies on phytochemical diversity often use Shannon’s diversity index, calculated as

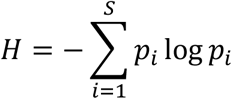

where *S* is the total number of compounds in the sample and *p*_*i*_ is the relative abundance (proportion) of compound *i*. This index takes evenness into account, such that for a given number of compounds, diversity is maximized when they occur at equal proportions. For diversity measures also considering disparity, functional diversity indices such as Rao’s Q can be used. For phytochemical diversity, Rao’s Q measure the average dissimilarity between two randomly drawn compounds, weighted by their abundance, from a sample. It is calculated as

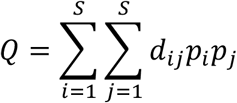

where *p*_*i*_ and *p*_*j*_ are the relative abundances of compounds *i* and *j*, and *d*_*ij*_ is the dissimilarity between compounds *i* and *j*. In this way, a dissimilarity matrix containing pairwise dissimilarities between phytochemical compounds, calculated based on biosynthetic or structural properties of the molecules (Box 2), can be included in measures of phytochemical diversity. While these traditional diversity indices are frequently used, a consensus has developed that Hill numbers represent a more suitable way of quantifying diversity (Ellison, 2010). Hill numbers, also referred to as Hill diversity or effective number of species (Hill, 1973; Jost, 2006; Chao *et al*., 2014), are related to the traditional indices, and defined as

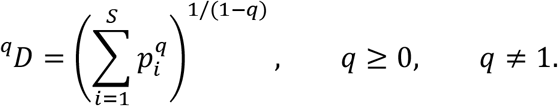

This measure is undefined for *q* = 1, but this can still be calculated because its limit as *q* approaches 1 equals

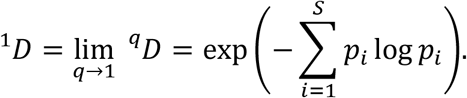

The parameter *q* is the diversity order, and controls the sensitivity of the measure to the relative abundances of the compounds. For *q* = 0, the measure is simply equal to the number of compounds, so that ^*0*^*D = S*. For *q* = 1, compounds are weighed in proportion to their abundance, and ^*1*^*D* is equal to the exponential of Shannon’s diversity. For *q* > 1, more weight is put on abundant compounds, and at *q* = 2, ^*2*^*D* is equal to the inverse Simpson diversity. Using Hill numbers to measure diversity has several advantages (Chao *et al*., 2014). First, the parameter *q* controls the sensitivity of the measure to the relative abundances of compounds. Adjusting *q*, the behaviour of the index can be controlled to enable a more nuanced measure of diversity. Second, Hill numbers are expressed in units of effective numbers, which is the number of equally abundant compounds required to obtain the same value of diversity. In this way, the units behave intuitively, facilitating comparisons between groups. Third, partitions of Hill numbers into α-, β-and γ-diversity is straightforward (Jost, 2007). Finally, Hill numbers can be generalized to a measure of functional diversity, so that compound dissimilarity can also be taken into account (Chiu & Chao, 2014; Chao *et al*., 2014). In this way, it is possible to measure several types of functional diversity in the Hill numbers framework. The most central of these is (total) functional diversity, which can be calculated as

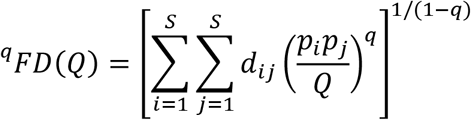

where *Q* is Rao’s Q (Chiu & Chao, 2014). This measure is also undefined for *q* = 1, but its limit as *q* approaches 1 equals

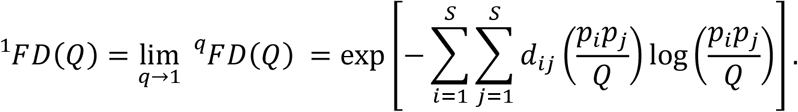

The index ^*q*^*FD(Q)* is a function of all three diversity components. This functional diversity quantifies the effective total dissimilarity between compounds in a sample (Chiu & Chao, 2014). It can therefore be used as a comprehensive measure of phytochemical diversity, sensitive to variation in richness, evenness and disparity. Overall, Hill numbers provide a unified approach to quantifying phytochemical diversity but so far only the non-functional version (^*q*^*D*) has been used in a few studies (e.g. Marion *et al*., 2015; Cosmo *et al*., 2021; Philbin *et al*., 2021). Diversity measures combining richness, evenness and disparity into a single metric may obscure independent variation in each component. However, Hill numbers enable separate and combined quantification of all three components. As mentioned, Hill diversity at *q* = 0 simply equals the richness, while at *q* = 1 it is dependent on richness and evenness. Functional Hill diversity adds a layer of data by also considering disparity. At *q* = 0, it is equal to the sum of the pair-wise dissimilarities in the dissimilarity matrix, a measure known as functional attribute diversity (Walker *et al*., 1999). At *q* = 1, it is a measure sensitive to all three components of diversity. For a given number of compounds, functional Hill diversity increases with increasing compound dissimilarities, and, in contrast to Rao’s Q (Shimatani, 2001), is always maximised at complete evenness. Evenness can also be calculated in this framework (Tuomisto, 2012). Thus, the Hill numbers framework can quantify all components of diversity. Overall, it is crucial to understand the indices’ behaviour, and additional ways of calculating diversity exist (Petchey & Gaston, 2006; Chao *et al*., 2019). Hill numbers measure α-diversity, quantifying the diversity within a single sampling unit. Quantifying differences between samples can be done by calculating β-diversity from measures of α-and γ-diversity (Jost, 2007). Alternatively, and more common in chemical ecology, Bray-Curtis dissimilarities can be calculated between samples, and visualized with a non-metric multidimensional scaling (NMDS) plot (Brückner & Heethoff, 2017). Bray-Curtis dissimilarities measure the compositional dissimilarity between samples, but do not take compound dissimilarity into account. A method to do so was developed by Junker (2018), who calculated a biosynthetically informed dissimilarity measure using Generalized UniFrac dissimilarities (Chen *et al*., 2012). Here, compound dissimilarities are calculated based on the proportion of shared enzymes, which is then incorporated in calculations of sample dissimilarities as Generalized UniFrac dissimilarities, such that two samples containing more biosynthetically different compounds have a higher dissimilarity. Collectively, with Hill numbers and Generalized UniFracs, it is possible to quantify both phytochemical diversity within sampling units and phytochemical dissimilarity between sampling units in a way that considers compound dissimilarities (Fig. 1). Therefore a generalized way to quantify compound dissimilarities is needed (Box 2).

**Fig. 1.**
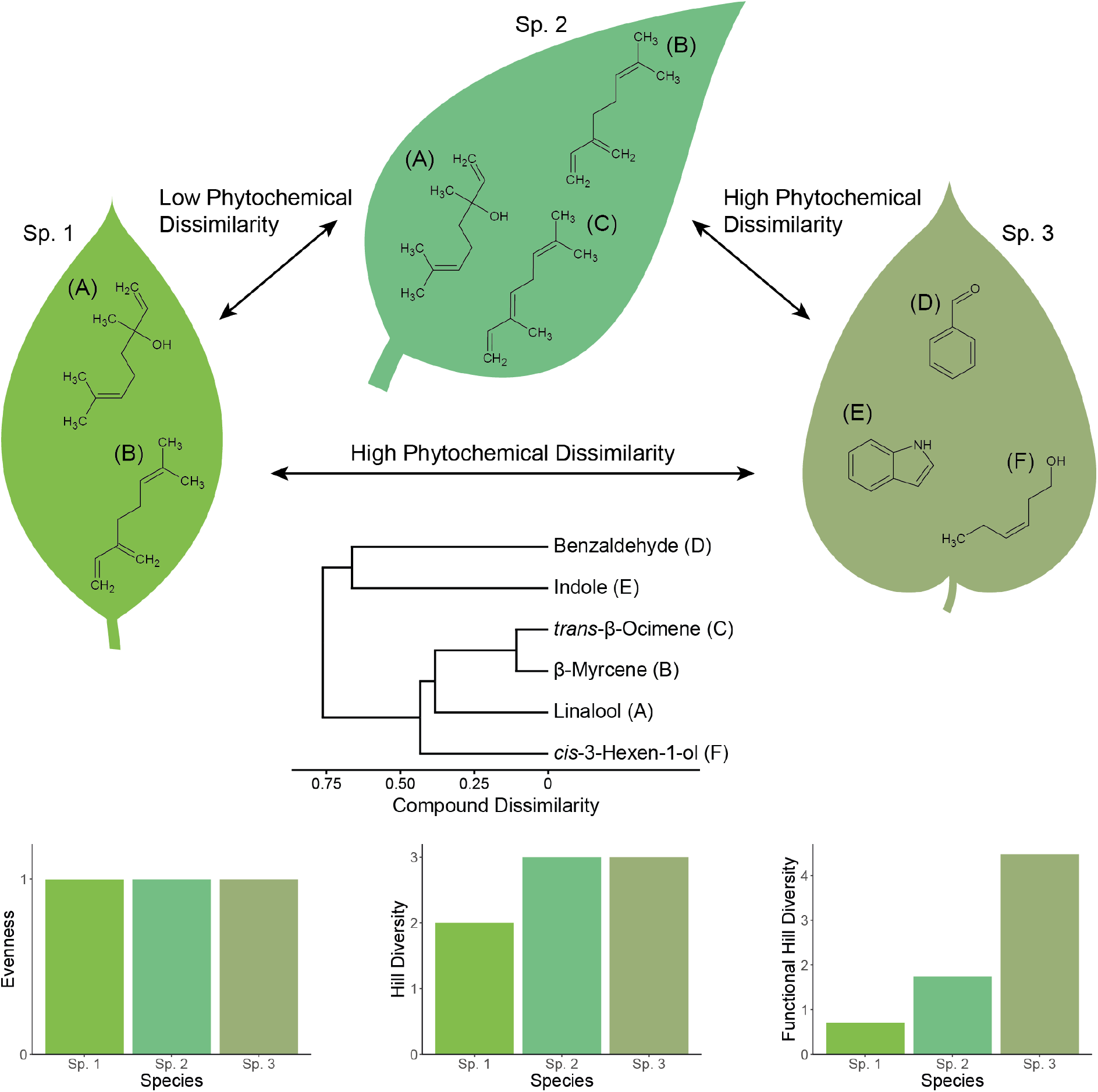
A conceptual illustration of how phytochemical diversity and dissimilarity is quantified. Leaves from three different plant species contain different phytochemicals, at equal abundances. Species 1 and 2 contain two and three structurally similar monoterpenes (linalool, β-myrcene, *trans*-β-ocimene), respectively. Species 3 contain three structurally more dissimilar compounds produced in different biosynthetic pathways (indole, an alkaloid; *cis*-3-hexen-1-ol, an aliphatic/fatty acid derivative; benzaldehyde, a benzenoid). The dendrogram illustrates structural dissimilarities between the compounds (calculated using *PubChem Fingerprints*). Species 1 and 2 contain similar compounds, and have a low phytochemical dissimilarity. Species 3 contains different compounds, and has a high phytochemical dissimilarity to the other species. The phytochemical diversity of the species depends on how it is quantified, indicated by the bar plots. All species have equal evenness. Hill diversity is lowest in species 1 because it contains only two compounds. Functional Hill diversity, taking compound dissimilarities into account, is higher in species 3 than in species 2, as an effect of the former having a set of more dissimilar phytochemicals.

### Box 2. Quantifying compound dissimilarities

Calculating functional diversity in the Hill numbers framework, and dissimilarity with Generalized UniFracs, requires a way to quantify dissimilarities between phytochemical compounds. We utilize three different complementary methods to quantify compound dissimilarity that only require knowing compound identities. The first method is based on a hierarchical classification of phytochemicals. H. W. Kim *et al*. (2021) developed *NPClassifier*, a deep-learning tool that, based on expert knowledge, automatically classifies natural products into three hierarchical levels (pathway, superclass and class) that largely correspond to the biosynthetic pathways the compounds are produced in. Using a similar approach as in Junker (2018), we use this classification to calculate Jaccard dissimilarities between phytochemicals, as a measure of their biosynthetic dissimilarity. The second method uses molecular fingerprints to quantify compound dissimilarities based on structural properties of the molecules (Cereto-Massagué *et al*., 2015). We use the *PubChem Fingerprint*, which consists of 881 binary variables representing the presence or absence of different features in the molecule, including specific elements, bonds and ring structures (Bolton *et al*., 2008; Cereto-Massagué *et al*., 2015). The fingerprints are then used to calculate Jaccard dissimilarities between compounds, as a measure of their structural dissimilarity. The third method is a graph-based *flexible Maximum Common Substructure* (*fMCS*) method (Cao *et al*., 2008b; Wang *et al*., 2013). The *fMCS* of two compounds is the largest substructure that occurs in both of them, allowing for a set number atom/bond mismatches in the identified substructures. By comparing the number of atoms in the common substructure to the total number of atoms in the molecules, Jaccard dissimilarities can be calculated based on *fMCS*, as a measure of their structural dissimilarity. Using *fMCS* is more computationally intensive than *PubChem Fingerprints*, but may have increased performance (Wang *et al*., 2013). Both measures have recently been successfully used to create mixtures of phytochemical compounds with different levels of structural diversity (Whitehead *et al*., 2021a). Using three different methods to quantify compound dissimilarity provides a choice upon which properties (biosynthetic: *NPClassifier*; structural: *PubChem Fingerprints, fMCS*) to compare phytochemicals (see Results and Discussion). Data needed for dissimilarity calculations is accessed by the *NPClassifier* tool (H. W. Kim *et al*., 2021), and the PubChem database (S. Kim *et al*., 2021) via functions in the *chemodiv* package.

## Methods

*chemodiv* is available as an R package on the Comprehensive R Archive Network, CRAN (https://CRAN.R-project.org/package=chemodiv). It contains a number of functions to easily calculate and visualize different types of phytochemical diversity and dissimilarity. The package utilizes other packages for retrieving and processing chemical data, including *webchem* (Szöcs *et al*., 2020), *ChemmineR* (Cao *et al*., 2008a) and *fmcsR* (Wang *et al*., 2013); and for diversity and dissimilarity calculations, including *vegan* (Oksanen *et al*., 2022), *hillR* (Li, 2018) and *GUniFrac* (Chen *et al*., 2022). In this section, we describe the functions of the package, and provide examples of analyses on real and simulated datasets. Details on calculations of diversity and quantification of compound dissimilarity are described in Box 1 and Box 2, respectively, and are jointly summarized in Fig. 1.

### Data requirements

Two sets of data are required to fully utilize the functions in the *chemodiv* package. First, a data set on the relative abundances of phytochemical compounds in different samples, as commonly obtained from GC-MS and LC-MS analyses. Second, a list with the common name, SMILES and InChIKey for all the compounds in the first dataset is needed. SMILES and InChIKey are chemical identifiers, and are readily compiled by searching for compounds in chemical databases such as PubChem (S. Kim *et al*., 2021), or using its automated tool Identifier Exchange Service. These identifiers are used to download data on biosynthetic and structural properties of the phytochemical compounds from different databases.

### Description of functions in the R package

The *chemodiv* package functions are summarized in Table 1. A full analysis of the diversity and dissimilarity of a set of phytochemical samples includes a number of largely sequential steps. First, the function *chemoDivCheck* can be used to check that datasets are correctly formatted. Second, the function *NPCTable* enables the use of the *NPClassifier* tool (H. W. Kim *et al*., 2021) directly within R, to classify compounds into three hierarchical levels largely corresponding to biosynthetic pathways. Third, the function *compDis* uses the list of compounds with their chemical identifiers to generate a dissimilarity matrix with dissimilarities between compounds, calculated based on the biosynthetic classification by *NPClassifier*, and/or structural properties of the compounds (*PubChem fingerprints, fMCS*; Box 2). Fourth, three different functions can be used to calculate different types of diversity for the samples. Function *calcDiv* calculates diversity within samples using the most common indices of α-diversity and evenness, including Shannon’s diversity, inverse Simpson diversity, Rao’s Q, two types of evenness, and both types of Hill diversity (Box 1). Functional Hill diversity and Rao’s Q use the dissimilarity matrix generated by *compDis* in the diversity calculations. Function *calcDivProf* can be used to generate a diversity profile, where both types of Hill diversity are calculated for a range of *q*-values. When plotted, a diversity profile can provide a more nuanced view of the diversity. Function *calcBetaDiv* calculates β-diversity as both types of Hill diversity. Fifth, the function *sampDis* generates a dissimilarity matrix with phytochemical dissimilarities between samples, calculating either Bray-Curtis or Generalized UniFrac dissimilarities, the latter of which uses the compound dissimilarity matrix generated by *compDis*. Sixth, functions *molNet* and *molNetPlot* generate and plot molecular networks, where nodes represent compounds and edges (links) represent similarities between compounds. Such networks can illustrate dissimilarities between compounds, calculated by *compDis*, and simultaneously visualize their abundances. Finally, the function *chemoDivPlot* can be used to conveniently create basic plots of the calculated measures of compound dissimilarity, sample diversity and sample dissimilarity, for different groups of samples that may represent treatments, populations, species or similar. Additionally, the function *quickChemoDiv* is a shortcut function that uses the other functions to calculate or visualize phytochemical diversity for a dataset in a single step. The central parts of the workflow are shown in Fig. 2. A detailed demonstration of the functions is included in a vignette in the package. All functions produce output in standard formats, facilitating statistical tests on the diversity and dissimilarity measures.

**Table 1.** Overview of the functions available in the *chemodiv* package, and a description of what they do.

| Function call with default parameters | Description |
| --- | --- |
| `chemoDivCheck(sampleData, compoundData)` | Checks that datasets are appropriately formatted |
| `NPCTable(compoundData)` | Classifies phytochemical compounds into a hierarchy of biosynthetic groups using *NPClassifier* |
| `compDis(compoundData, type = "PubChemFingerprint", npcTable = NULL, unknownCompoundsMean = FALSE)` | Calculates dissimilarities between compounds using biosynthetic (*NPClassifier*) and/or structural (*PubChem Fingerprints*, *fMCS*) properties |
| `calcDiv(sampleData, compDisMat = NULL, type = "HillDiv", q = 1)` | Calculates selected types of α-diversity and evenness measures, in both traditional and Hill numbers frameworks |
| `calcDivProf(sampleData, compDisMat = NULL, type = "HillDiv", qMin = 0, qMax = 3, step = 0.1)` | Calculates diversity profiles in the Hill numbers framework |
| `calcBetaDiv(sampleData, compDisMat = NULL, type = "HillDiv", q = 1)` | Calculates β-diversity in the Hill numbers framework |
| `sampDis(sampleData, compDisMat = NULL, type = "BrayCurtis", alpha = 1)` | Calculates Bray-Curtis and/or Generalized UniFrac dissimilarities between samples |
| `molNet(compDisMat, npcTable = NULL, cutOff = "median")` | Creates a molecular network based on compound dissimilarities, and calculates some network properties |
| `molNetPlot(sampleData, networkObject, groupData = NULL, npcTable = NULL, plotNames = FALSE, layout = "kk")` | Uses the output of *molNet* to create a plot of the molecular network |
| `chemoDivPlot(compDisMat = NULL, divData = NULL, divProfData = NULL, sampDisMat = NULL, groupData = NULL)` | Creates selected plots to visualize phytochemical diversity, compound dissimilarity and sample dissimilarity |
| `quickChemoDiv(sampleData, compoundData = NULL, groupData = NULL, outputType = "plots")` | Makes use of other functions in the package to calculate or visualize phytochemical diversity for a dataset in a single step |

**Fig. 2.**
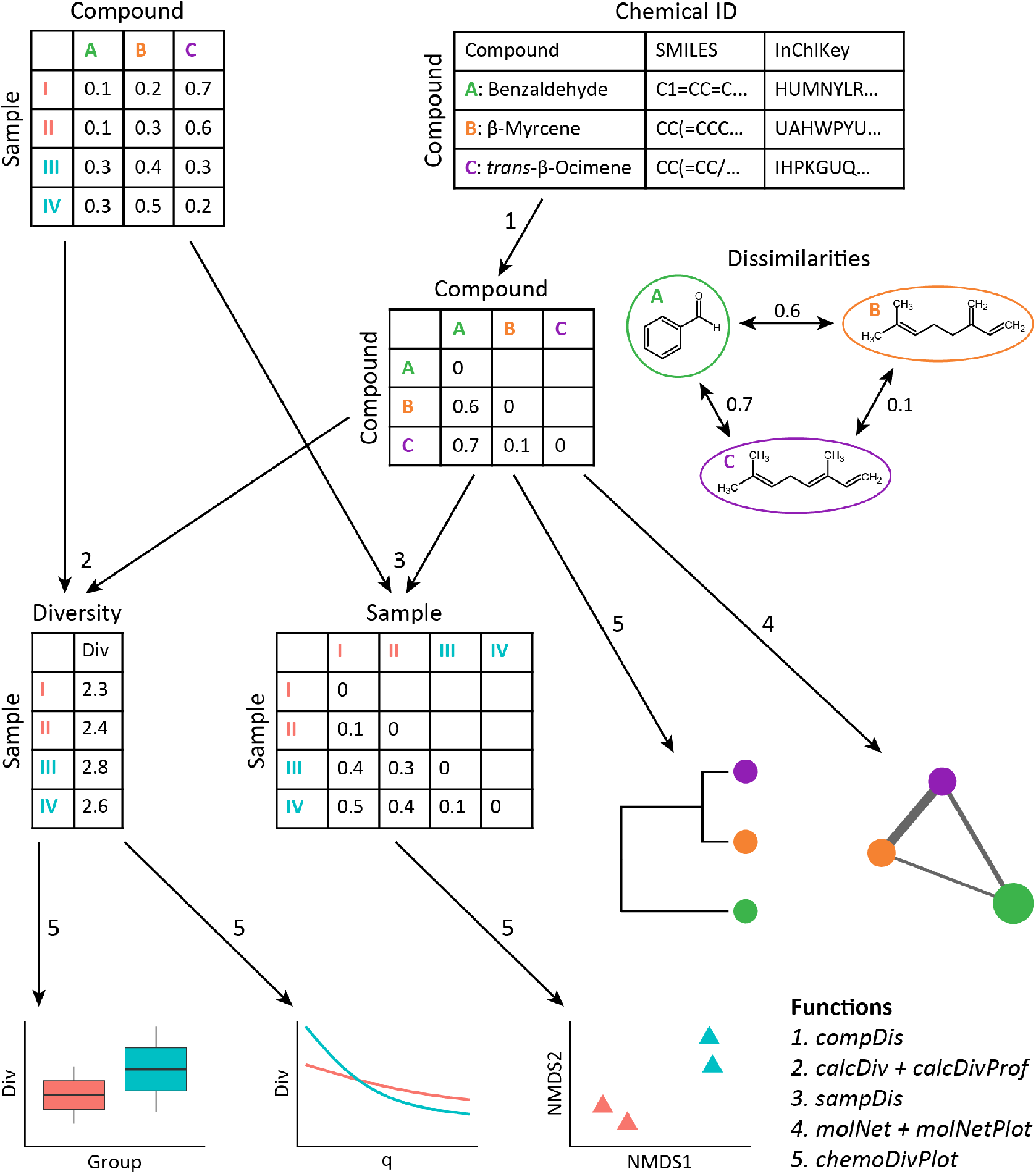
An illustration of the workflow of the main functions in the *chemodiv* package. A dataset with relative abundances of phytochemical compounds in four samples belonging to two different groups (red and blue), and a list with the common name, SMILES and InChIKey for the compounds in the dataset are required. The *compDis* function uses the list of compounds to generate a dissimilarity matrix with dissimilarities between compounds (1). β-Myrcene (B) and *trans*-β-ocimene (C) are two linear monoterpenes that have a low structural dissimilarity, while benzaldehyde (A), a benzenoid, is more dissimilar to the other compounds. In combination with the sample dataset, the compound dissimilarity matrix is used to calculate phytochemical diversity within samples (2; functions *calcDiv* and *calcDivProf*) and phytochemical dissimilarity between samples (3; function *sampDis*). Functions *molNet* and *molNetPlot* are used to create a molecular network (4), while *chemoDivPlot* is used to create multiple plots of compound dissimilarity, sample diversity and sample dissimilarity (5).

### Examples on simulated and real datasets

To demonstrate the applicability of the *chemodiv* package for measuring phytochemical diversity and dissimilarity, we analysed a number of simulated and real datasets with it.

The first example includes a semi-simulated dataset on plant defence compounds. Glucosinolates is a class of phytochemicals produced by most species in the Brassicaceae family. They provide protection against generalist herbivores (after being hydrolysed by myrosinase), but are less efficient against specialist herbivores (Hopkins *et al*., 2009). Plants in the *Erysimum* (Brassicaceae) genus have additionally gained a novel chemical defence in the form of cardenolides, a class of phytochemicals that provides protection against some glucosinolate-adapted specialist herbivores. A diverse mixture of phytochemicals from both groups could therefore maximise herbivore protection (Züst *et al*., 2018, 2020). However, quantifying such diversity is not straightforward, as e.g. Shannon’s diversity do not consider that these two groups of compounds contain structurally different molecules produced in different biosynthetic pathways. To demonstrate the applicability of calculating measures of phytochemical diversity and dissimilarity that take such difference into account, we used a haphazardly selected set of eight glucosinolates and eight cardenolides found in *Erysimum cheiranthoides* L. (Züst *et al*., 2020; Mirzaei *et al*., 2020). Thereafter, by sampling from different normal distributions, we simulated data on the relative concentration of these compounds in three groups of 16 individual plants each: (1) plants with a high concentration of glucosinolates and a low concentration of cardenolides, (2) plants with a low concentration of glucosinolates and a high concentration of cardenolides, and (3) plants with a high concentration of four glucosinolates and four cardenolides, and a low concentration of the remaining glucosinolates and cardenolides. Using the functions in the package, we quantified compound dissimilarity based on the structure of the compounds using *fMCS*, calculated functional Hill diversity and a corresponding diversity profile, calculated sample dissimilarities using Generalized UniFracs, and visualized results.

The second example is a fully simulated dataset designed to further examine the use of functional Hill diversity as a measure of phytochemical diversity. This example includes a base dataset with relative concentrations of eight phytochemicals (four at high concentration, four at low concentration) simulated in a similar way as example one. This time, also compound dissimilarities were simulated, by sampling from a binomial distribution. We then created additional groups of samples that had an increased richness, evenness and/or compound dissimilarity for a total of eight groups with all combinations of high and low values for the three diversity components. We then calculated and plotted compound richness, evenness and both versions of Hill diversity for the different groups.

The third example is a dataset on floral scent from Larue *et al*. (2016), where floral volatiles were collected from *Achillea millefolium* L. and *Cirsium arvense* L. in a scent manipulation experiment. Here we only include plants in the control treatment. Using the package functions, we calculated and created plots of the compound dissimilarities (based on *PubChem Fingerprints*), functional Hill diversity and sample dissimilarities (using Generalized UniFracs), and created molecular networks to compare the two species.

Finally, we compared the three methods (*NPClassifier, PubChem fingerprints* and *fMCS*) for quantifying compound dissimilarities. For this, we used a collection of 2 855 phytochemical compounds from the KEGG database (Kanehisa & Goto, 2000). The heaviest 20% of molecules were excluded to reduce computation times. Then, from this subset, 20-40 compounds were randomly selected, and dissimilarity matrices were calculated using the *compDis* function with the three different methods. Mantel tests were then used to calculate correlation coefficients between matrices. This was repeated 50 times, and results were plotted in order to examine how comparable compound dissimilarities generated with the different methods were to each other.

## Results and Discussion

### Evaluating examples on simulated data

Analyses for the semi-simulated dataset with cardenolides and glucosinolates exemplify how the structural component of phytochemical diversity can be quantified. Compound dissimilarity, quantified using *fMCS*, is low among glucosinolates and among cardenolides, but higher when comparing glucosinolates to cardenolides, as evident by the dendrogram separating the two groups of compounds (Fig. 3a). These differences in compound dissimilarity influence the phytochemical diversity measured as functional Hill diversity (Fig. 3b). Even without variation in compound richness or evenness, there are clear differences in the diversity of samples from the different groups. Diversity is lowest for the group with high concentration of only glucosinolates, intermediate for the group with high concentration of only cardenolides (due to a somewhat higher average compound dissimilarity among cardenolides than among glucosinolates) and highest for the group containing a high concentration of compounds from both classes. The diversity profile displays functional Hill diversity for *q* = 0 to *q* = 3, varying how much weight is put on low-concentration compounds (Fig. 3c). At *q* = 1 (also shown in Fig. 3b), equal weight is put on all compounds. At *q* = 0, compound proportions are not taken into account and the functional Hill diversity is thus equal for all three groups. Lastly, the NMDS illustrates Generalized UniFrac dissimilarities of samples (Fig. 3d). Samples cluster in groups, as an effect of being dominated by different compounds. Within-group dispersion is highest for the group containing high concentrations of both types of compounds, as a result of higher average compound dissimilarity. A comparison between the diversities and dissimilarities calculated here, and the traditionally used Shannon’s diversity and Bray-Curtis dissimilarities is shown in Fig. S1.

**Fig. 3.**
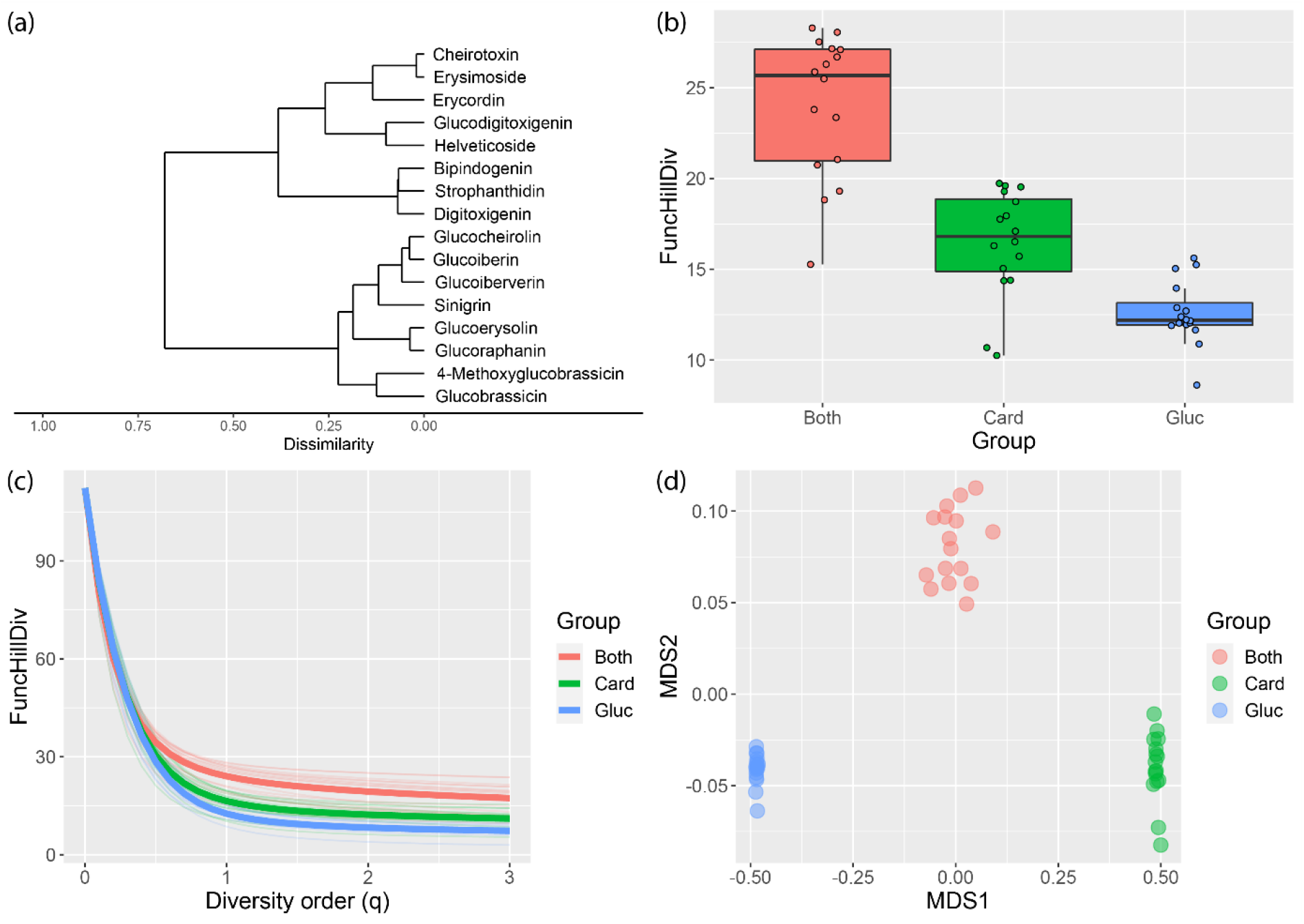
Phytochemical diversity and dissimilarity for the semi-simulated dataset with glucosinolates and cardenolides, visualized by the *chemoDivPlot* function in the *chemodiv* package. (a) Dendrogram of compound dissimilarities based on *fMCS*, with a clear separation between cardenolides (upper branch) and glucosinolates (lower branch). (b) Functional Hill diversity (*q* = 1) for the groups containing a high concentration of cardenolides (Card), a high concentration of glucosinolates (Gluc), and both (Both). (c) Diversity profile showing the functional Hill diversity for *q* = 0 to *q* = 3. Thick lines represent group means while thin lines represent individual samples. (d) NMDS plot visualizing sample dissimilarities (Generalized UniFracs) between the three groups.

Results from the second example, with the fully simulated dataset, are summarized in Fig. S2. In short, this example illustrates the behaviour of different diversity measures, and demonstrates the overall suitability of using functional Hill diversity as a measure of phytochemical diversity. By simulating samples with low and high richness, evenness and compound dissimilarity, as expected we found that functional Hill diversity is lowest when all three components have low values, intermediate when some components have high values and other have low values, and highest when all three components have high values.

While many studies have found that phytochemical diversity, measured as e.g. Shannon’s diversity, can shape interactions between plants and other organisms (e.g. Iason *et al*., 2005; Glassmire *et al*., 2016; Tewes *et al*., 2018), the structural dimension of phytochemical diversity may also be important for ecological interactions (Richards *et al*., 2015; Junker *et al*., 2018; Cosmo *et al*., 2021). In the example with glucosinolates and cardenolides, the group with structurally dissimilar compounds from two different biosynthetic pathways had the highest diversity when measured as functional Hill diversity. On a general level, two structurally similar molecules can be expected to have a more similar biological activity than two structurally dissimilar molecules (Berenbaum & Zangerl, 1996; Martin *et al*., 2002), although there are contrasting examples where e.g. different enantiomers have different function (He *et al*., 2019). Therefore, a set of structurally dissimilar phytochemicals from different biosynthetic pathways may be more diverse in regards to its function (Philbin *et al*., 2022), with potential effects on plant fitness. For example, increased structural diversity of phytochemicals in leaves, quantified from ^1^H-NMR spectra, has been found to decrease herbivory in multiple *Piper* species (Glassmire *et al*., 2019; Cosmo *et al*., 2021; Philbin *et al*., 2022). Additionally, Whitehead *et al*. (2021a) found that increasing the structural diversity of phenolics in the diet of eight insect and fungi plant consumers increased the proportion of those consumers negatively affected by the phenolics. Hence, calculations of compound dissimilarity based on molecular structure (*PubChem Fingerprints, fMCS*), and subsequent measures of sample diversity or dissimilarity, may inform about how diverse in regards to function a set of phytochemical compounds is (Berenbaum & Zangerl, 1996; Philbin *et al*., 2022).

If compound dissimilarities are instead calculated based on *NPClassifier*, this may help to account for non-independence of compounds due to shared biosynthetic pathways (Junker, 2018). Additionally, this informs about biosynthesis differences between sets of compounds, which can be useful for studies on the evolution of these pathways and their importance for producing different types of compounds for herbivore protection (Becerra *et al*., 2009). While efficient, this method has a lower resolution compared to using manually collected data on enzymes (Junker, 2018), because the classification is limited to three hierarchical levels. It should also be noted that structural dissimilarity has been used as a proxy for biosynthetic similarity in other studies (Dowell & Mason, 2020; Cna’ani *et al*., 2021), and in our simulations dissimilarities calculated with all three methods were correlated (Fig. S3), indicative of an overall consistency between methods. Additionally, the structural and biosynthetic similarity of compounds may correlate with similarity of physicochemical properties such as volatility, reactivity and polarity, that may be ecologically important (Rasmann & Agrawal, 2011; Conchou *et al*., 2019). Researchers should make a deliberate choice of how to quantify compound dissimilarities based on what questions are addressed. Overall, the structural and biosynthetic components of the compounds, readily quantified by the *chemodiv* package, are important parts of the phytochemical diversity that should be included in measures of it.

### Evaluating examples on real data

Analyses of the *A. millefolium* and *C. arvense* dataset indicate that phytochemical diversity of the floral scent bouquet was higher in the latter species (Fig. 4b-c), mainly due to a higher average number of compounds (*A. millefolium* = 36.3, *C. arvense* = 48.4). The floral scent composition was also clearly different between species (Fig. 4d). Illustrations of compound similarities by the molecular networks (Fig. 5), indicate the presence of two main clusters of structurally similar compounds mainly consisting of the pathways “Terpenoids” and “Shikimates and Phenylpropanoids” respectively. The scent bouquet of *A. millefolium* plants was dominated by compounds from the first group (Fig. 5a), while the scent bouquet of *C. arvense* plants was dominated by compounds from the second group (Fig. 5b).

**Fig. 4.**
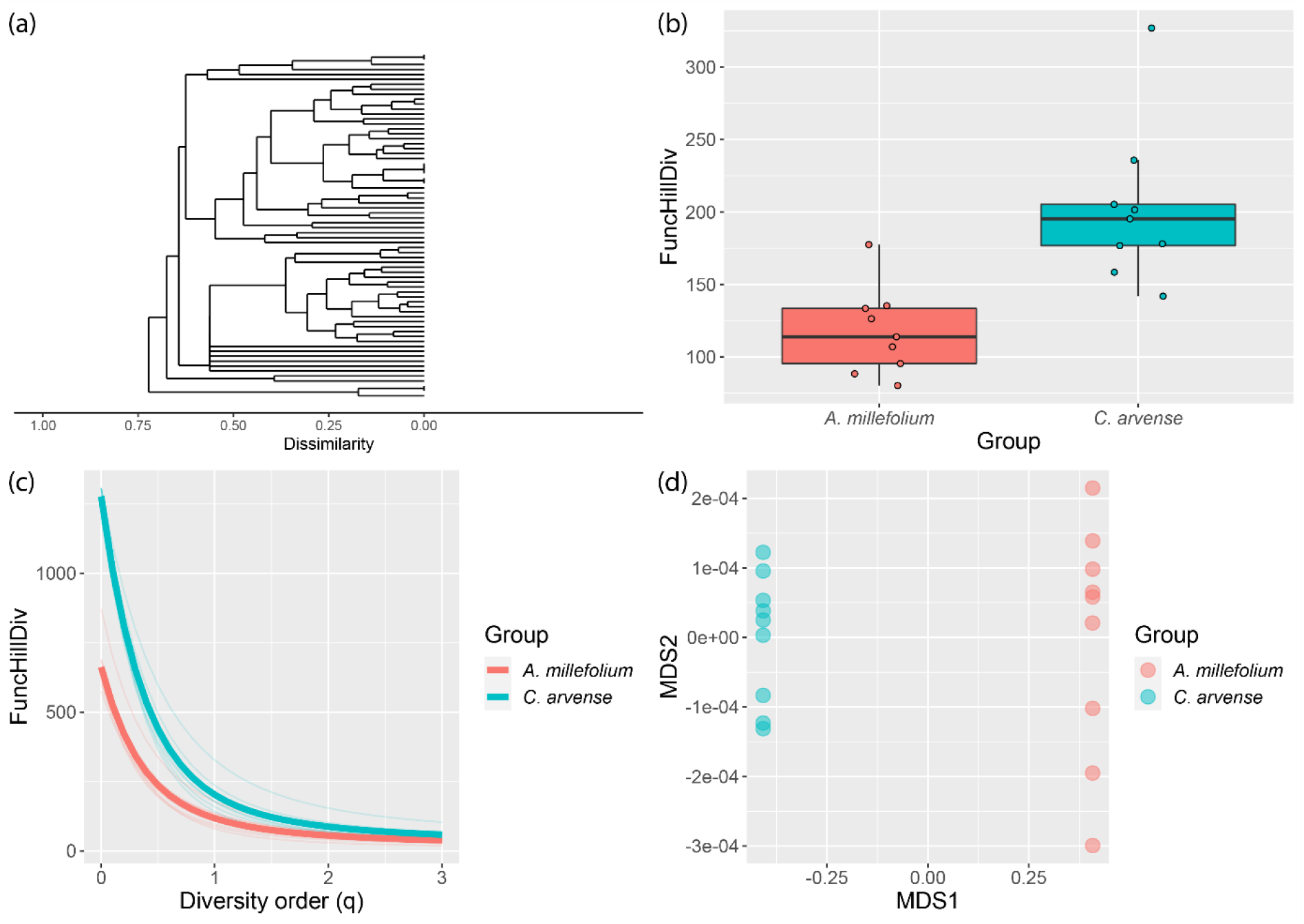
Phytochemical diversity and dissimilarity for *Achillea millefolium* and *Cirsium arvense* (*n* = 9 for both species), visualized by the *chemoDivPlot* function in the *chemodiv* package. (a) Dendrogram of compound dissimilarities based on *PubChem Fingerprints* (compound names have been excluded for clarity). (b) Functional Hill diversity (*q* = 1) for the two species. (c) Diversity profile showing the functional Hill diversity for *q* = 0 to *q* = 3. Thick lines represent species means while thin lines represent individual samples. (d) NMDS plot visualizing sample dissimilarities (Generalized UniFracs) between the species.

**Fig. 5.**
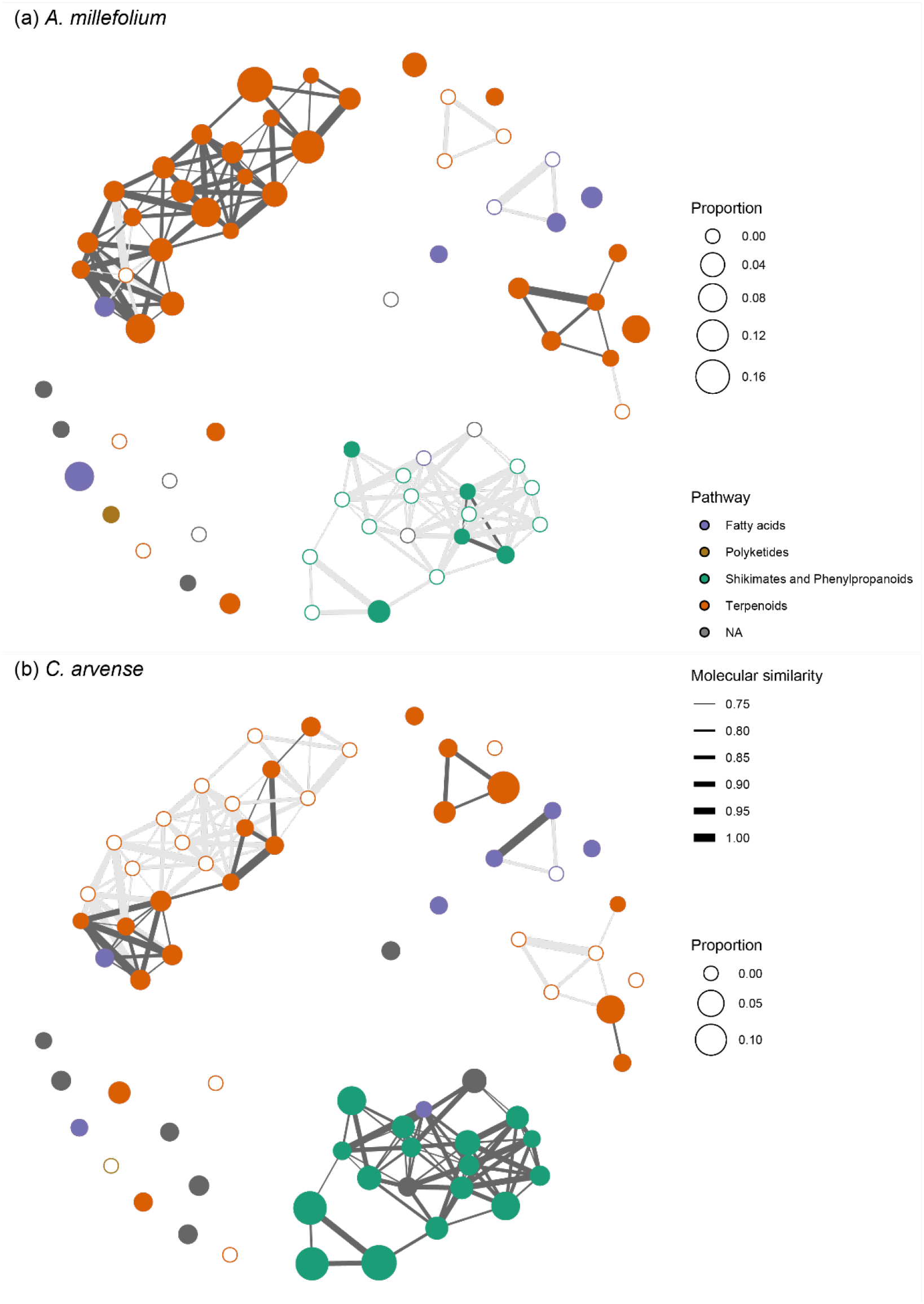
Molecular networks of the compounds found in *Achillea millefolium* (a) and *Cirsium arvense* (b), visualized by the *molNetPlot* function in the *chemodiv* package. Edge width represents similarities between compounds (with a cut-off of 0.75) and node colour represents the pathway classification from *NPClassifier*, indicating to which major biosynthetic group compounds belong. These are identical in (a) and (b). Node size represents proportional concentration (mean values for each species), and differs between (a) and (b). Note that nodes with white fill represent zero values, i.e. compounds not present in that species, and that edges connecting to such nodes are a lighter shade of grey.

Most examples on the effect of phytochemical diversity on ecological interactions regard herbivores, where the diversity represents a complex phenotype important for herbivore defence through toxic effects of compounds during consumption (Marion *et al*., 2015; Kessler & Kalske, 2018). In contrast, for a pollinator in search of nectar, or a herbivore searching for host plants, phytochemical diversity, in the form of volatile organic compounds (VOCs), represents information in a complex environment (Kessler, 2015; O’Connor *et al*., 2019). With potential correlations between compound properties and neural/behavioural response (Khan *et al*., 2007; Haddad *et al*., 2008; but see Knaden *et al*., 2012), a structurally diverse set of VOCs may be functionally diverse, and may therefore enable a generalist plant to efficiently attract different pollinators and/or simultaneously repel antagonistic insects (Schiestl, 2010; Junker & Blüthgen, 2010; Gershenzon *et al*., 2012; Junker, 2016). In other cases, diverse mixtures of leaf VOCs can make it difficult for herbivores to locate suitable host plants Zu *et al*. (2020). Using comprehensive measures of diversity may enable a better understanding of its effects on both antagonistic and mutualistic interactions between plants and other organisms.

### Applicability and caveats

*chemodiv* allows chemical ecologist to easily and comprehensively analyse phytochemical diversity. A few aspects should be considered when using the package and quantifying diversity. The package requires that compounds have been identified in order to quantify compound dissimilarity. The functions can handle unknown compounds, but *chemodiv* is of most use for datasets where most compounds are identified. If that is not the case, diversity can instead be calculated with indices not considering compound dissimilarities. Alternatively, other methods to quantify the dissimilarity of unknown compounds can be used. For example, in metabolomic analyses where individual compounds are not identified, the GNPS ecosystem provides methods for calculating cosine similarities between MS/MS spectra (Wang *et al*., 2016), which could be used in diversity calculations.

Even if chemodiversity is an ecologically relevant measure, other aspects of the phytochemical phenotype are undoubtedly also important. First, functions such as pollinator attraction, herbivore defence or plant-bacteria interactions often depend on individual compounds rather than mixtures (Junker, 2016; Zhou *et al*., 2017; Burdon *et al*., 2018). However, such examples often regard the effect of a compound on a single interacting species, while in nature, plants simultaneously interact with numerous mutualists and antagonists. Each of these might select for the occurrence of different compounds, resulting in evolution of increased chemical diversity (Berenbaum & Zangerl, 1996; Iason *et al*., 2011; Whitehead *et al*., 2021a). Second, an important aspect of the phytochemical phenotype is total abundance, which of course may be important for function. Diversity indices consider relative values, such that two samples with the same compounds in identical proportions will have identical diversity, even if they differ in absolute concentrations. Related to this, the number of compounds detected in a sample may partly depend on total abundance (Wetzel & Whitehead, 2020). Therefore, direct comparisons of phytochemical diversity should ideally be done for samples collected with identical methods. If this is not the case and total abundances vary substantially among samples, diversity can be quantified using Hill numbers at higher diversity orders (e.g. *q* = 2). Doing so, less weight is put on low concentration compounds, which may be less functionally important, decreasing any potential influence of differences in total abundance on measures of diversity.

Notably, quantification of molecular structural dissimilarity is not limited to phytochemicals. Diversity of chemical mixtures have been quantified for e.g. fungi VOCs (Guo *et al*., 2021), snake venom (Holding *et al*., 2021), coralline algae metabolites (Jorissen, 2021) and fish fatty acids (Feiner *et al*., 2018). Instead of using traditional measures, such studies can also include the structural dissimilarities of compounds for more comprehensive measures of chemodiversity. It is also worth mentioning that we have focused on quantifying diversity on the level of individual samples, often likely to represent individual plants. Others have instead utilized phytochemical compounds as functional traits to measure diversity on a community level (Salazar *et al*., 2016), representing a complementary approach which can answer questions related to trait-based ecology and niche processes (Müller & Junker, 2022; Walker *et al*., 2022).

### Conclusions

Plants produce a remarkable number of phytochemicals. By now, it is widely accepted that, rather than being metabolic waste products, they are functionally important, and their diversity is the result of adaptive processes (Hartmann, 2007). However, much is still unknown about how the complex phenotype that is the composition of phytochemical compounds affects interactions between plants and other organisms (Wetzel & Whitehead, 2020). We believe the diversity of the compounds, including their structural and biosynthetic properties, to be an important dimension of this variation that deserves further attention. The *chemodiv* package provides an easy yet comprehensive way to quantify this diversity for many types of data collected by chemical ecologists. By providing this tool, we hope to give researchers the opportunity to more efficiently test in what ways phytochemical variation influences ecological interactions and evolutionary processes, which should increase our understanding of the vast diversity of phytochemical compounds found in plants.

## Acknowledgements

We thank the members of the Research Unit FOR 3000 “Ecology and Evolution of Intraspecific Chemodiversity of Plants” for fruitful discussions on the topic of chemodiversity, and members of the Junker lab for helpful comments on the manuscript. This project was funded by the German Research Foundation (Deutsche Forschungsgemeinschaft, DFG, JU 2856/5-1).

## Author Contributions

All authors conceived the study and contributed to study design. HP evaluated statistical methods and created the R package with input from RRJ. HP wrote the manuscript with contributions from RRJ and TGK. All authors approved of the final version of the manuscript.

## Data Availability

The R package is available on CRAN (https://CRAN.R-project.org/package=chemodiv) and developed on GitHub (https://github.com/hpetren/chemodiv). Scripts and data of the examples in the paper will be uploaded to a public data repository upon acceptance of the paper for publication.

## Supporting Information

**Fig. S1.**
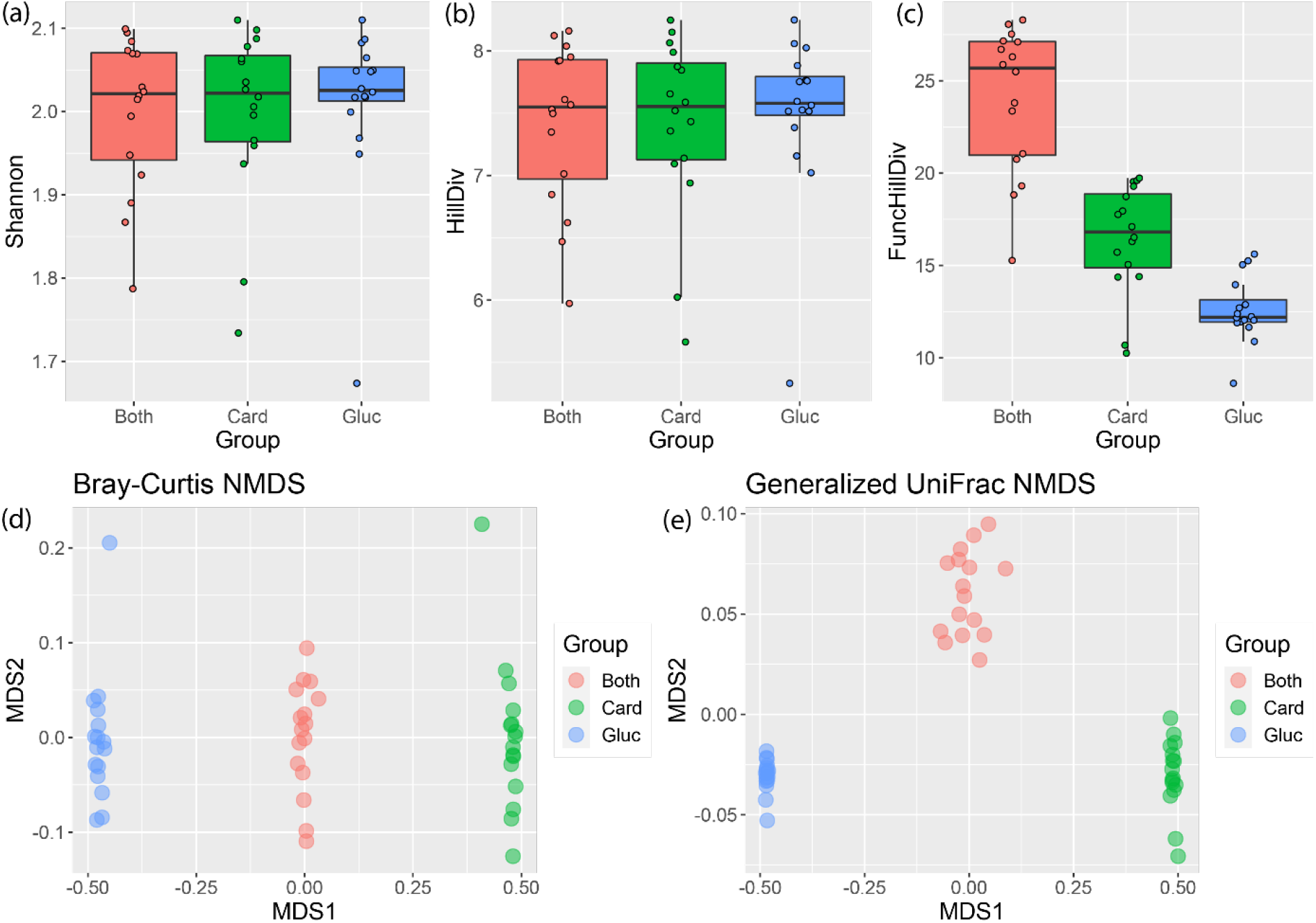
Comparison of different measures of phytochemical diversity (a-c) and dissimilarity (d-e) for the semi-simulated dataset with cardenolides and glucosinolates. For measures of phytochemical diversity, Shannon’s diversity (a) and Hill diversity (b) (equal to the exponential of Shannon’s diversity) do not take compound dissimilarity into account, and all three groups have similar diversity. In contrast, functional Hill diversity (c) depends also on compound dissimilarity, with the result that the group with a high concentration of both cardenolides and glucosinolates (red) has a higher phytochemical diversity than the groups with only cardenolides (green) or only glucosinolates (blue) at high concentrations. For measures of phytochemical dissimilarity, when Bray-Curtis dissimilarities are used, the within-group dispersion is similar in all three groups. In contrast, when Generalized UniFrac dissimilarities are used, which take compound dissimilarity into account, the group with both types of compounds at a high concentration has a higher within-group dispersion, as an effect of containing a mixture of more dissimilar compounds.

**Fig. S2.**
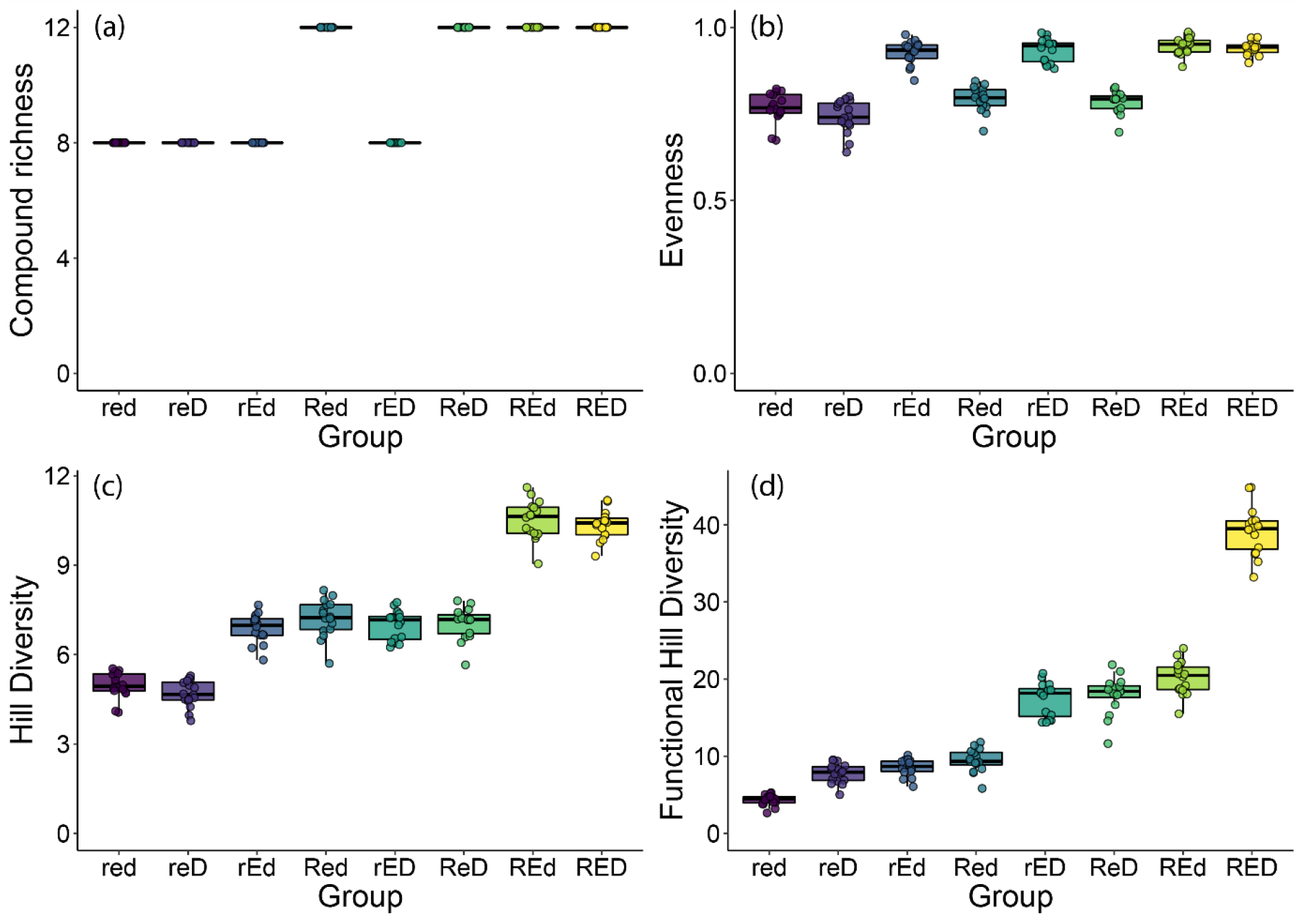
Compound richness (a), Pielou’s evenness (b), Hill diversity (c) and functional Hill diversity (d) for eight different groups of simulated phytochemical samples. Each group consists of 16 samples simulated to have a high or low richness (r/R), evenness (e/E) and compound dissimilarity (d/D). Lowercase letters indicate a low value, uppercase letters indicate a high value. Richness (a) is the number of compounds in the samples (and is equal to Hill diversity at *q* = 0). Evenness (b) depends only on the relative abundances of compounds. Hill diversity (*q* = 1) (c), equal to the exponential of Shannon’s diversity, depends on both richness and evenness, and is therefore higher for groups with high richness and/or evenness. Functional Hill diversity (*q* = 1) (d) is dependent on all three components of diversity (richness, evenness and disparity). It is lowest when richness, evenness and disparity is low, intermediate when one or two of the components is high, and highest when richness, evenness and dissimilarity are all high. In this regard, functional Hill diversity is therefore the most comprehensive measure of diversity.

**Fig. S3.**
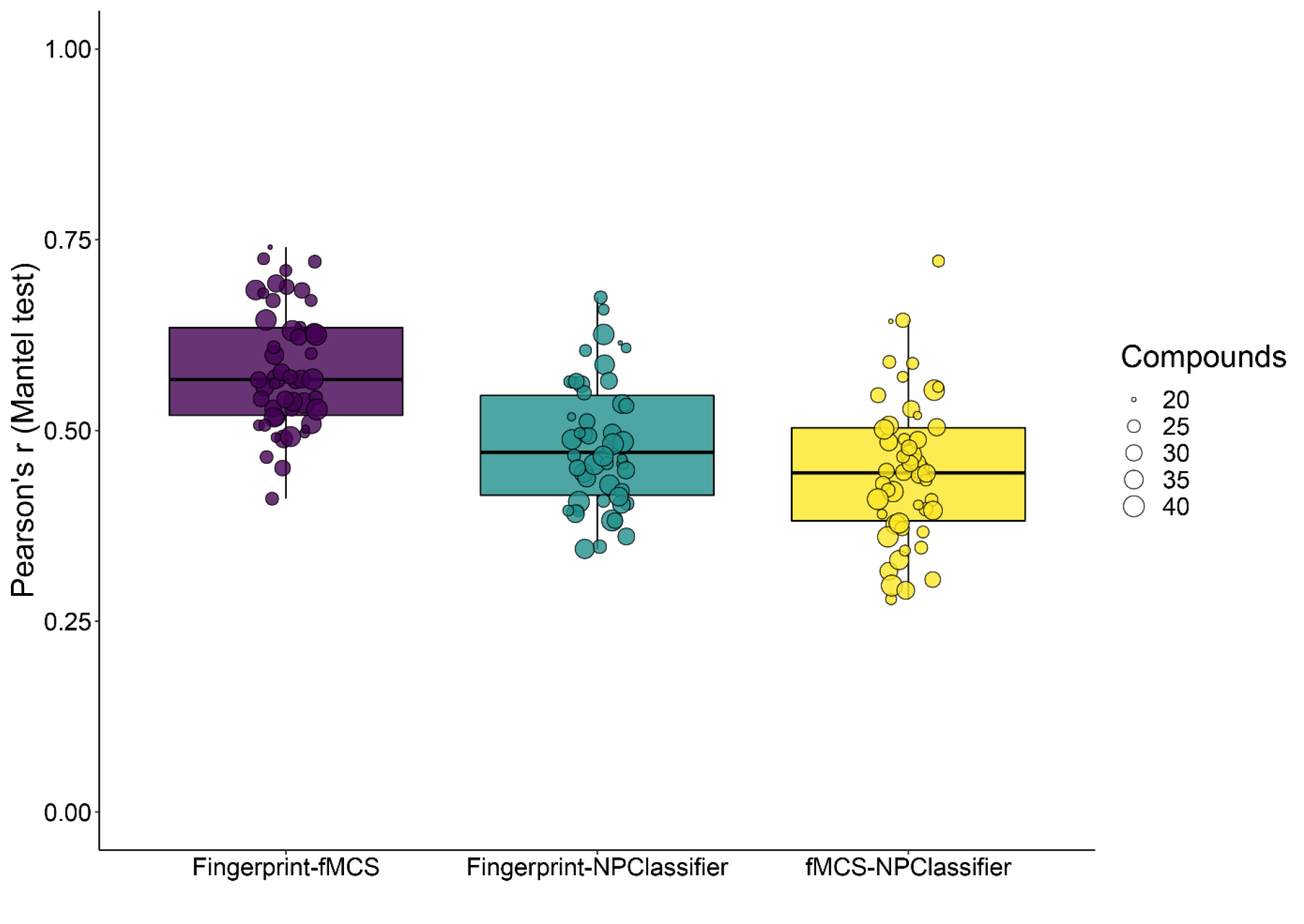
Comparison of compound dissimilarities calculated using the three different methods in the *compDis* function. In 50 iterations, 20-40 phytochemical compounds were randomly selected, and compound dissimilarities were calculated using three methods: *NPClassifier*, which compares compounds based on a classification of compounds into groups largely corresponding to biosynthetic pathways, and *PubChem fingerprints* and *fMCS*, which compare compounds based on structural properties of the molecules using binary fingerprints and substructure matching, respectively. Mantel tests were then used to calculate Pearson’s correlation coefficients between dissimilarity matrices. Each box includes the median, and upper and lower quartiles of these correlation coefficients. Data points of individual comparisons are overlaid. Correlations were overall relatively strong, and statistically significant (*P* < 0.05) in all cases. Mean correlation coefficients were highest between dissimilarity matrices based on *PubChem fingerprints* and *fMCS* (mean *r* = 0.58), and somewhat lower between dissimilarity matrices based on *PubChem fingerprints* and *NPClassifier* (mean *r* = 0.48) and dissimilarity matrices based on *fMCS* and *NPClassifier* (mean *r* = 0.45).

